## Supplementary figures and images for "Protective role of protease-activated receptor-2 in anaphylaxis model mice"

### Figure S1

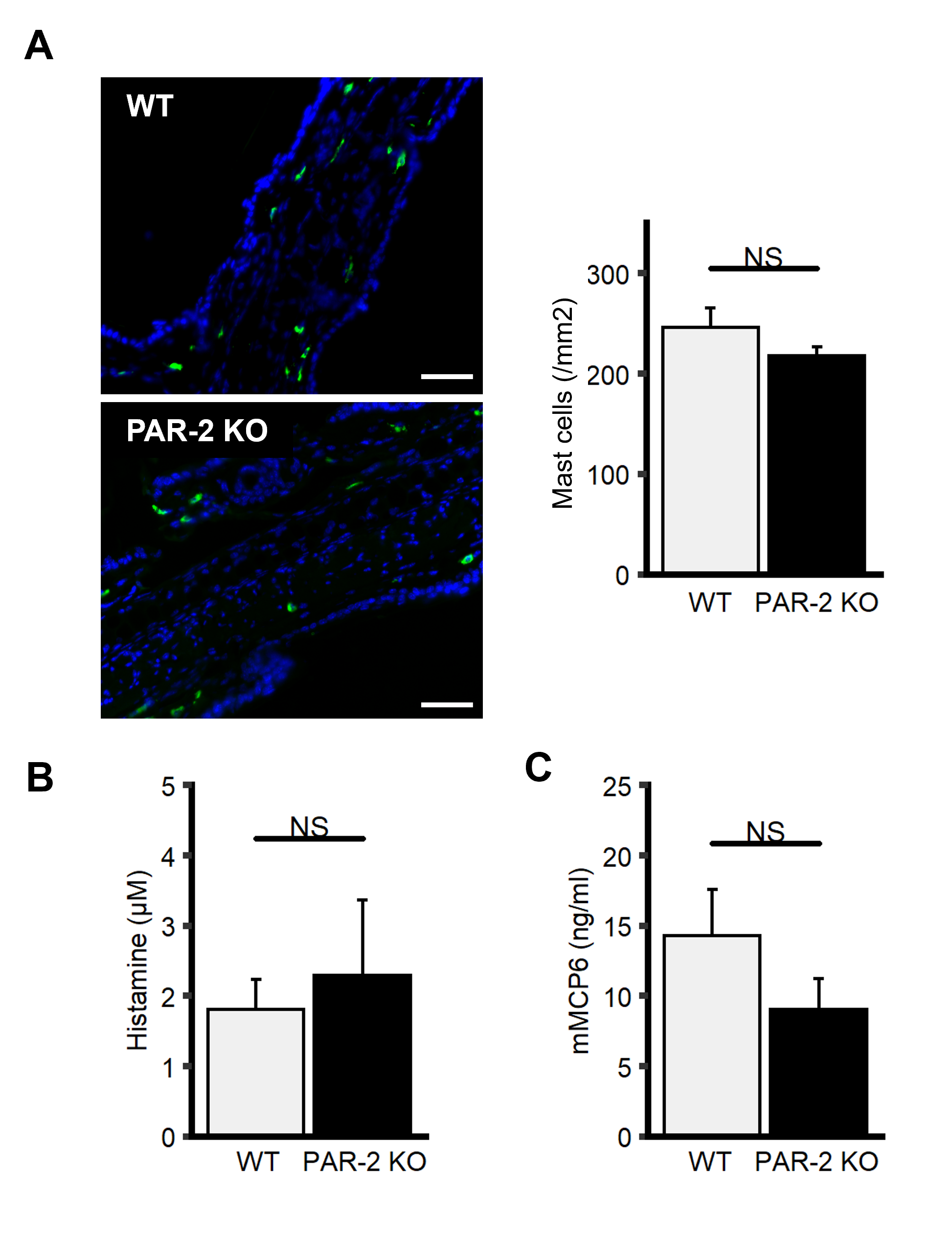

### Figure S2

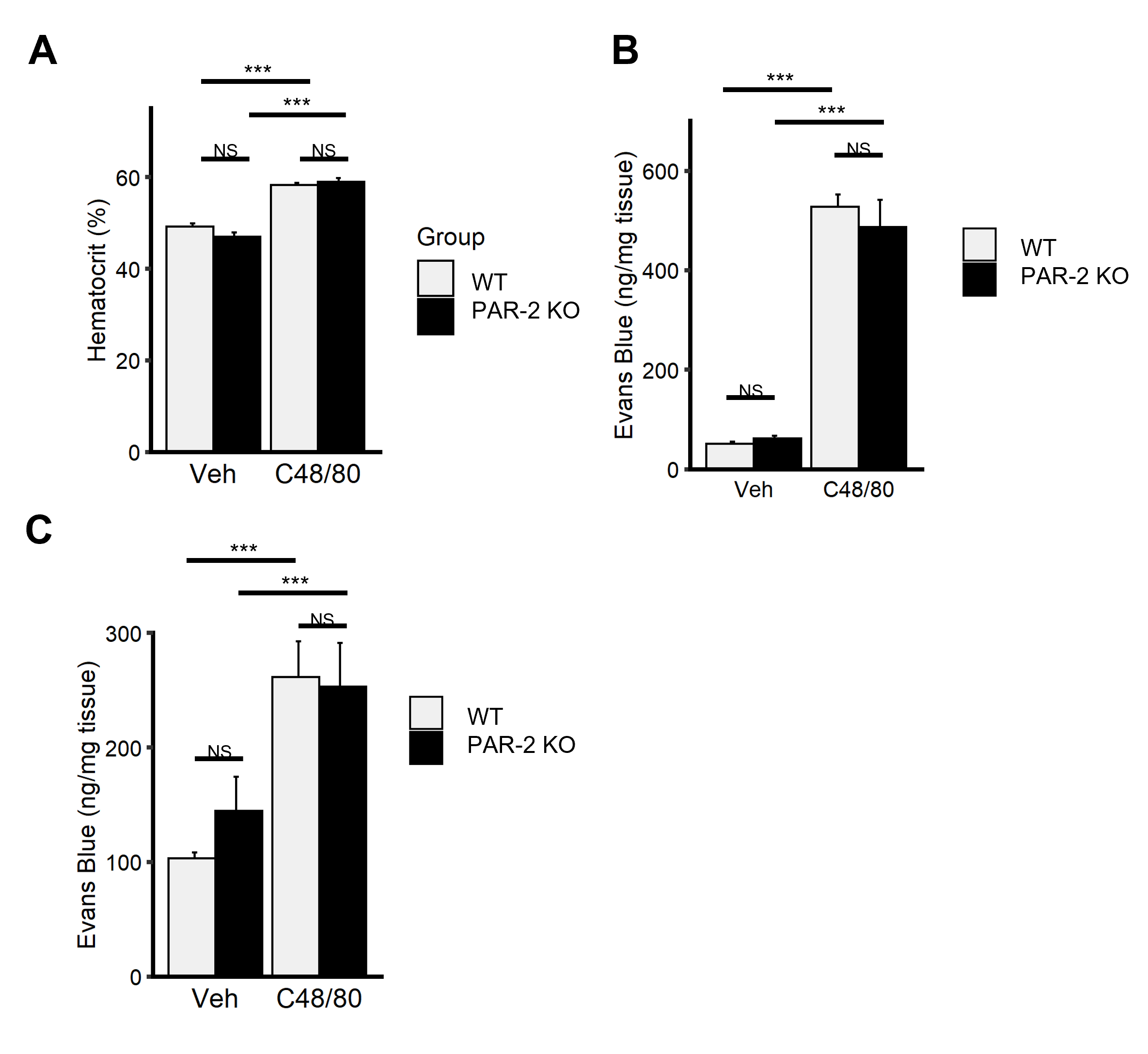
